## Supplementary Infomration File for "Heterozygous missense variant in *GLI2* impairs human endocrine pancreas development"

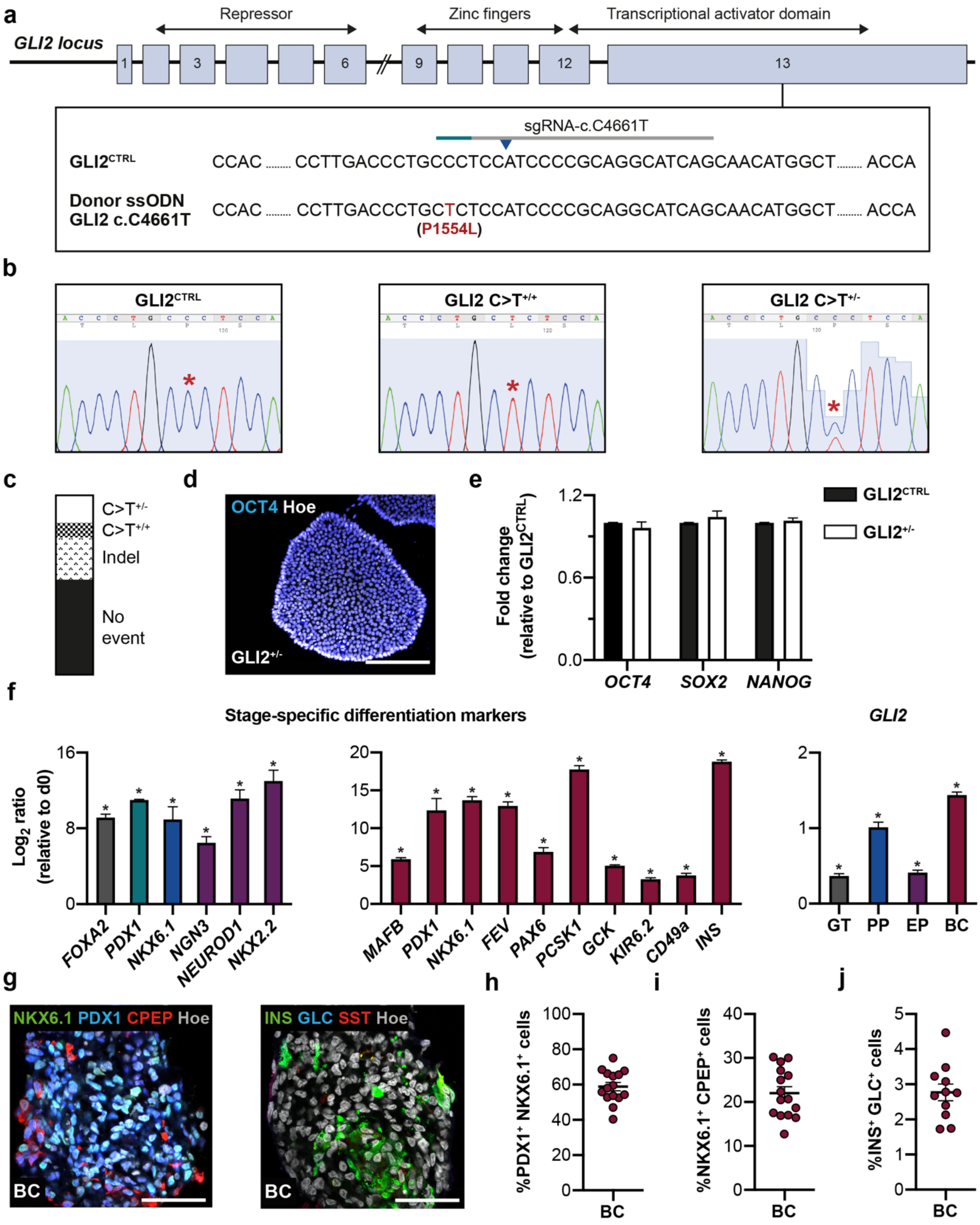


**Supplementary Figure 1. Generation of patient-like GLI2 c.C4661T iPSC lines.**

**a** CRISPR gRNA design for generating GLI2 c.C4661T disease variant using the HMGUi001-A2 iPSC line. The target sequence of the sgRNA and the protospacer-adjacent motif (PAM) sequences are indicated in grey and green, respectively. The mutation was introduced through homology direct repair using a ssODN template (Supplementary Table 2). The blue arrow indicates the predicted Cas9 cleavage site.

**b** Sequencing-graphs of WT control line ($GLI2^{CTRL}$) and heterozygous ($GLI2^{+/-}$) and homozygous ($GLI2^{+/+}$) c.C4661T mutant clones. Red asterisk indicates C>T switch.

**c** CRISPR Cas9 efficiency. We established several heterozygous (14.71% efficiency) and homozygous (8.82% efficiency) c.C4661T mutant iPSC clones, as confirmed by Sanger sequencing.

**d** Alkaline phosphatase (AP) staining of representative $GLI2^{+/-}$ iPSC clone.

**e** RT-qPCR analysis of selected pluripotency gene transcripts in WT control line ($GLI2^{CTRL}$) and CRISPR-Cas9 engineered $GLI2^{+/-}$ iPSC line. Values are normalized to GAPDH and relative to WT iPSCs. Values shown are mean ± SEM. n=3.

**f** RT-qPCR analyses of stage-specific markers (left) and *GLI2* (right) expression in iPSC-based differentiation model. Values are normalized to *GAPDH* and relative to day 0. Statistical significance calculated using two-tailed Student's t-test. *$P<0.05$.

**g** Representative immunofluorescence staining images of differentiated β-like cell (BC) clusters at day 21 for PDX1, NKX6.1, human C-PEPTIDE (CPEP) and INSULIN (INS), GLUCAGON (GLC), SOMATOSTATIN (SST). Nuclei were labeled with Hoechst. Scale bar, 20 µm.

**h** Scatter plot shows percentage (%) of cells double positive for PDX1 and NKX6.1 in differentiated BC clusters.

**i** Scatter plot shows percentage (%) of cells double positive for NKX6.1 and human C-PEPTIDE at day 21.

**j** Scatter plot shows percentage (%) INSULIN/GLUCAGON-double positive cells, representing polyhormonal cell fraction, at day 21. The number of positive cells in (**h, i, j**) staining was normalized to the total number of cells contained in each cluster and shown as %. Values shown are mean ± SEM.

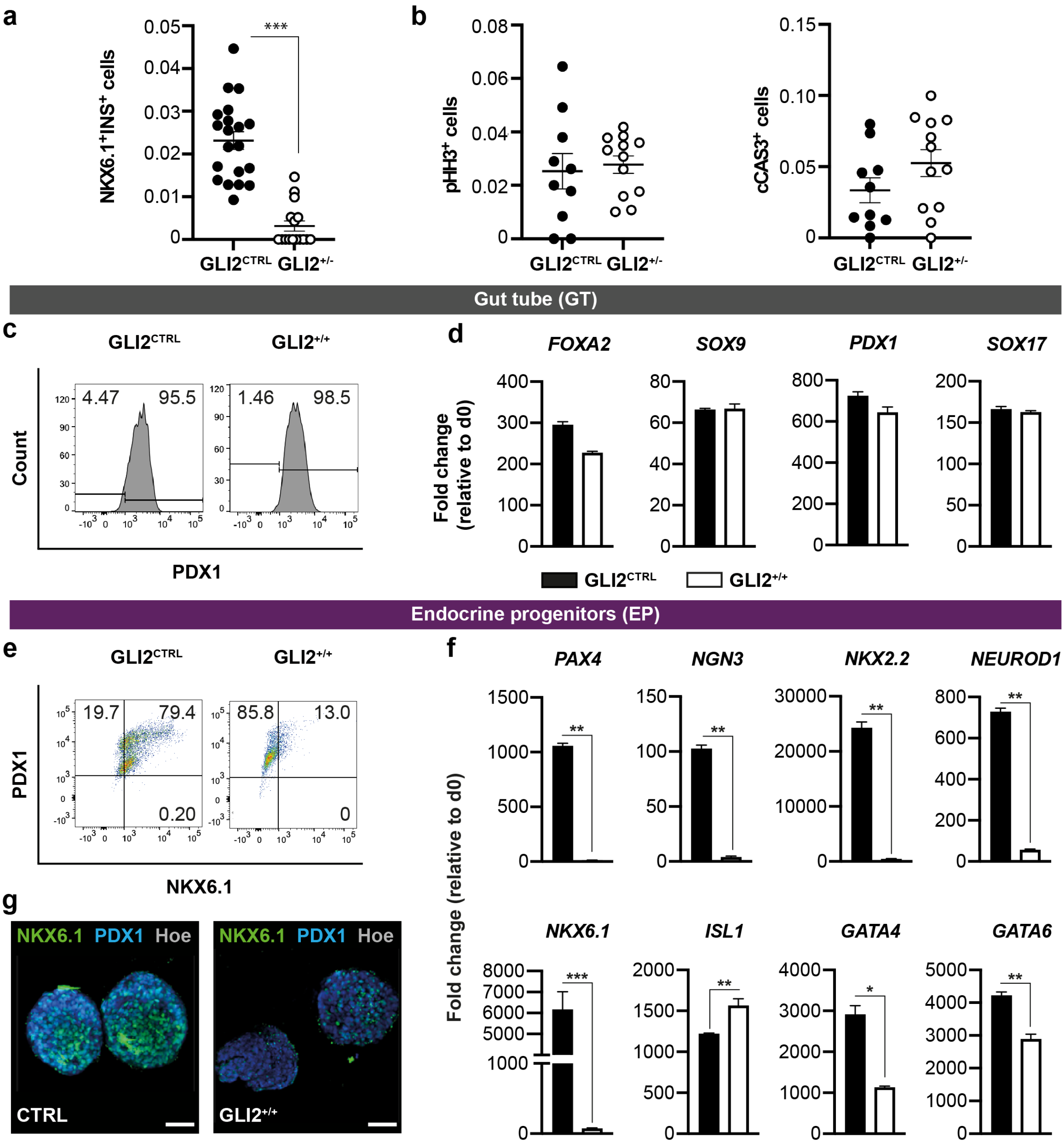


**Supplementary Figure 2. Characterization of homozygous GLI2$^{+/+}$ cells during pancreatic cell differentiation.**

**a** Scatter plot shows significant decrease of NKX6.1- and INSULIN-double positive cells in differentiated GLI2$^{+/-}$ clusters. The number of NKX6.1+/ INSULIN+ cells was normalized to the total number of cells contained in each cluster and shown as %. n = 3. ***p < 0.001; Student's t test.

**b** Quantification of the cells positive for the mitotic marker phosho-histone H3 (pHH3) and active caspase-3 (CAS3) in differentiated GLI2$^{+/-}$ clusters. n=3.

**c** Representative flow cytometry plots of PDX1$^{+}$ cells (shown as %) in GLI2$^{CTRL}$ and GLI2$^{+/+}$-derived cells at day (D) 5 of differentiation. n=3.

**d** RT-qPCR analysis of selected gene transcripts in GLI2$^{CTRL}$- and GLI2$^{+/-}$ differentiated cells at D5. Values are normalized to *GAPDH*. Data are shown as fold change relative to undifferentiated cells (d0). Values shown are mean ± SEM. n=3.
**e** Representative FACS plot of NKX6.1$^{+}$ and PDX1$^{+}$ cells (shown as %) in GLI2$^{CTRL}$- and GLI2$^{+/+}$-derived cells at D14. N=3.
**f** RT-qPCR analysis of selected gene transcripts in GLI2$^{CTRL}$ and GLI2$^{+/+}$ differentiated cells at D14. Data are represented as fold change relative to undifferentiated cells (d0). Values shown are mean ± SEM. n=2. *p <0.05; **p < 0.01, ***p < 0.001, Student's t test. The expression levels of *NGN3*, its downstream targets *NEUROD1*, *NKX2.2*, *PAX4*, were severely reduced in GLI2$^{+/+}$ mutant cells, while the expression of *ISL1* was upregulated.
**g** Whole-mount immunostaining for PDX1 and NKX6.1 in WT GLI2$^{CTRL}$ and GLI2$^{+/+}$-derived endocrine progenitor cells. Scale bars, 50 μm.

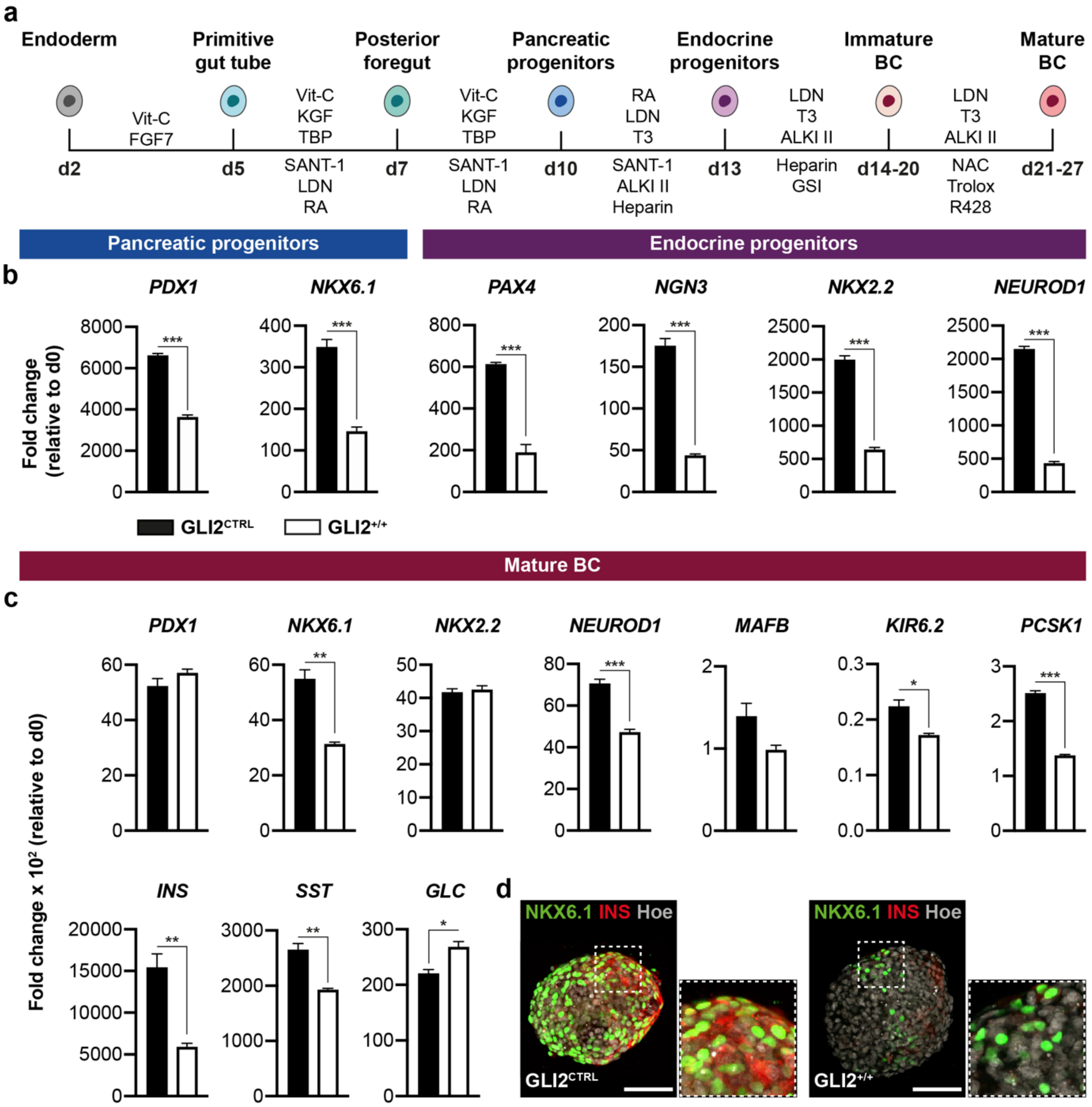


**Supplementary Figure 3. Directed differentiation of GLI2$^{+/+}$ iPSCs into β-like cells using an independent differentiation protocol[31].**

**a** Schematic diagram of the protocol[31] used to differentiate GLI2$^{CTRL}$ and GLI2$^{+/+}$ iPSCs into β-like cells (BC). For consistency with the previously employed protocol[29], we adapted the seven-stage protocol from Rezania et al.[31] to a 3D in suspension one and cells were differentiated on an orbital shaker.

**b, c** RT-qPCR analyses of indicated stage-specific markers. Values are normalized to GAPDH and relative to day 0. Values shown are mean ± SEM. n=3. *p <0.05; **p < 0.01, ***p < 0.001, Student's t test.

**d** Representative whole-mount immunostaining for NKX6.1 and INSULIN in $GLI2^{CTRL}$ and $GLI2^{+/-}$ iPSC-derived β-like cell clusters at day 27. Nuclei were labeled with Hoechst. Scale bars, 25 µm.

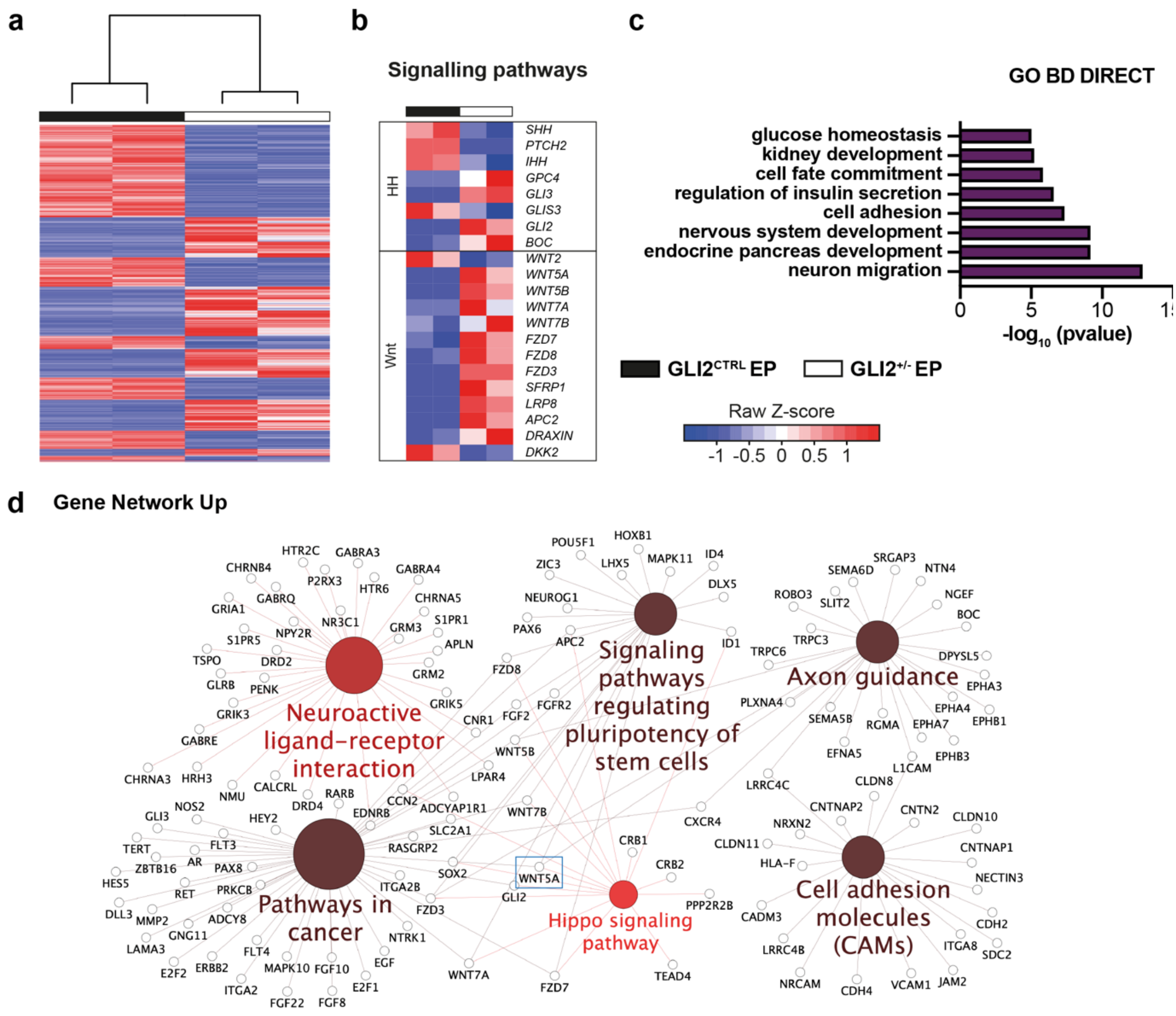


**Supplementary Figure 4. RNA-Seq analysis of GLI2$^{CTRL}$ and GLI2$^{+/-}$ iPSCs undergoing differentiation into endocrine progenitor cells.**

**a** Heatmap showing differentially expressed genes between GLI2$^{+/-}$- and GLI2$^{CTRL}$-derived EPs.

**b** Heatmap of selected differentially expressed genes between GLI2$^{+/-}$- and GLI2$^{CTRL}$-derived EPs. Boxes highlight genes belonging to the indicated signaling pathways. Colours represent high (red) or low (blue) expression values based on Z-score normalized to FPKM values for each gene.

**c** Gene ontology enrichment analysis of differentially expressed genes ($p<0.05$) performed on EP RNA-Seq datasets.

**d** GO term enrichment and pathway term network analysis of DEGs between GLI2$^{CTRL}$- and GLI2$^{+/-}$ -derived endocrine progenitors. Gene Network showing up-regulated genes. Term node size increases with term significance and colour corresponds to a particular functional group that is based on the similarity of their associated genes. Groups that shared 50% or more of the same genes were merged. The proportion of each node that is filled with colour reflects the kappa score. Functionally related groups partially overlap.

**Supplementary Table 1.** *In silico* pathogenicity predictions of the heterozygous *GLI2* p.P1554L variant^.

| Tool | Prediction |
|---|---|
| **SIFT**[1] | Deleterious |
| **PolyPhen2**[2] | Probably Damaging |
| **PROVEAN**[3] | Deleterious |
| **CONDEL**[4] | Deleterious |
| **CADD PHRED**[5] | 28.4 |

^The variant **rs767802807** has been reported in ClinVar (RCV002036029) as a variant of uncertain significance. Several heterozygous *GLI2* mutations have been reported to cause forebrain and pituitary defects.[19-21] However, these symptoms were not present in any of the sequenced patients.

[1]https://sift.bii.a-star.edu.sg/www/SIFT_seq_submit2.html;
[2]http://genetics.bwh.harvard.edu/pph2/;
[3]http://provean.jcvi.org/index.php;
[4]https://bbglab.irbbarcelona.org/fannsdb/help/condel.html;
[5]https://cadd.gs.washington.edu/.
